# Mannitol ingestion causes concentration-dependent, sex-specific mortality in adults of the fruit fly (*Drosophila melanogaster*)

**DOI:** 10.1101/564898

**Authors:** Katherine Fiocca, Meghan Barrett, Edward A. Waddell, Cheyenne McNair, Sean ’Donnell, Daniel R. Marenda

## Abstract

Mannitol, a sugar alcohol used in commercial food products, induced sex-specific mortality in the fruit fly *Drosophila melanogaster* when ingested at a single concentration (1M), and female mortality was greater than male mortality. We hypothesized that sex differences in energy needs, related to reproductive costs, contribute to increased mortality in females compared to males. To test for the effects of reproductive costs, we compared longevity to 21 days of actively mating and non-mating flies fed various concentrations of mannitol. We also asked whether mannitol-induced mortality was concentration-dependent for both males and females, and if mannitol’s sex-specific effects were consistent across concentrations. Females and males both showed concentration-dependent increases in mortality, but female mortality was consistently higher at all concentrations above 0.75M. Fly longevity to 21 days decreased further for both sexes when housed in mixed sex vials (as compared to single sex vials), suggesting the increased energetic demands of reproduction for both sexes may increase ingestion of mannitol. Mannitol fed to larvae did not alter emerging adult sex ratios, suggesting that sex-specific mortality due to mannitol occurs only in adults.

## Introduction

D-mannitol (henceforth mannitol) is a 6-carbon polyol produced via microbial fermentation, particularly by yeasts, and is the most common naturally-occurring polyol in plants and fungi [1-4]. Mannitol is commonly used as a sweet additive in consumer products as it is only partially absorbed in the human small intestine without increasing insulin secretion or blood glucose [1,5].

Mannitol produces a variety of gastrointestinal, reproductive, and survival effects when fed to other organisms [2,6-7]. Mannitol reduced survival and prevented adult female reproduction in *Pimpla turionellae* ichneumonid wasps [7]. In contrast, mannitol stimulated feeding behavior at low doses (72.6mM) in red flour beetles (*Tribolium castaneum*) and increased the longevity of females in comparison to males [8-9]. In contrast, adult female *Drosophila melanogaster* fed 1M mannitol food showed significantly decreased longevity over a seventeen-day trial in comparison to males [10].

We hypothesized that the sex-specific effects of mannitol in *D. melanogaster* could be caused by differing energetic demands between males and females. Oogenesis requires greater protein intake [11-12], leading females to eat more food, more frequently [13-14]. Sex-specific differences in survival between males and females could be due to differences in ingestion that increase self-dosing of mannitol in females. To test this hypothesis, first we assessed if mortality was concentration dependent in both males and females, suggesting dose-dependency, and if sex-specific differences in survival were consistent across concentrations. Next, we assessed if flies differed in survival when cultured in single sex vials or when cultured in vials with the opposite sex. Prior studies suggest female fly feeding rates increase substantially when housed, and mating, with males [15-17], so we predicted increased female mortality when cultured in mixed-sex vials. Reproduction can also be energetically costly for males due to courtship, competition, and mating behaviors, as well as sperm production, so we predicted increased male mortality in mixed-sex vials as well [18-22].

To test whether increased female mortality was inherently related to sex differences, such as genetic effects, we tested whether this sex-specificity also occurred during immature developmental stages. We reared larvae on mannitol foods of increasing concentration and found no difference in emerging adult sex ratios. Results suggest that sex-specific mortality is consistent across concentrations in adults only, and mannitol’s lethality is amplified when male and female flies are cultured together. Finally, we explore possible mechanisms of mannitol-induced mortality.

## Materials and Methods

### Culturing Drosophila

Wild-type (Canton S) *D. melanogaster* were raised to adulthood on standard *Drosophila* food for laboratory culturing and reared in an insect growth chamber at 27.5°C, 50% relative humidity, with a 12-h:12-h photoperiod [23]. Foods were prepared in 100 ml batches as follows: 9.4g cornmeal, 3.77g yeast, 0.71g agar, 0.75 ml Propionic acid, 1.88 ml Tegosept (10% w/v methyl p-hydroxybenzoate in 95% ethanol), 9.42 ml molasses. The treatment-specific amount of D-mannitol (HiMedia; GRM024-500G, Lot 000249743) was added, and beakers were filled with distilled water to a final volume of 100 ml. After heating the mixed ingredients to set the agar, foods were poured into vials and cooled until consistency was firm and uniform. An excess of food was provided, with 10 ml of food in each vial.

### Testing Concentration-Dependent Sex-Specific Adult Mortality in Single and Mixed-Sex Vials

Adult wild-type flies that were 0-24 hours post-eclosion were transferred from the standard food to the treatment foods. Two types of control food (no mannitol), consisting of the standard recipe with and without molasses, and treatment foods with mannitol concentrations of 0.25M, 0.5M, 0.75M, 1M and 2M were used. No molasses was used in the mannitol conditions. All foods contained 0.05% Brilliant Blue R-250 dye (used to visually check flies for food ingestion on day one of the trial). Vials contained ten flies each, with three vials of 10 female flies, three vials of 10 male flies, and three vials of 5 female and 5 male flies used per treatment (n=90 flies/treatment). Flies were moved to fresh foods of the same formulation every four days. Every 24 hours for 21 days, dead flies were removed from their vials, counted, and sexed. Dead flies were also dissected under a dissecting scope, to look for the presence of blue dye or other artefacts of mannitol ingestion in the gut. Photos of representative fly guts were taken on day four of mannitol ingestion from additional 0M and 0.75M treatment vials set up specifically for photos, after the initial round of dissections and data collection.

### Testing Effects of Feeding Mannitol to Larvae on Adult Sex Ratios and Eclosion Time

Groups of 15 male and 15 female wild-type flies raised on standard food were placed in vials containing 0M, 0.4M, or 0.8M mannitol adult foods and allowed to mate and lay for 24 hours before the adults were removed. Nine vials were used per concentration, with a total of 405 flies of each sex used for laying. Vials were checked for newly emerged adults every twelve hours from Day 10 to Day 15, and every twenty-four hours from Day 15 to Day 24 (when relatively few adults emerged). Adult flies were removed from the vials and sexed, and their eclosion day was recorded to the nearest 24 hours (0M: n=904; 0.4M: n=1264; 0.8M: n=262 adults).

### Statistical Analyses

Analyses were performed using SPSS v. 24 software, Graphpad Prism v. 8.0.0, and Sigmaplot v. 12.5 [24-26]. Sex-specific adult survival across concentrations were assessed using survival analyses in SPSS [27], with subjects living to the end of the trial or lost to reasons other than death (i.e., escaped or injured in handling) included in the analysis as right-censored values. Differences in survival distributions across concentrations, by housing condition (single vs. mixed sex), and by sex (for adults) were tested using pairwise log-rank Mantel Cox survival analysis tests. Mean pr(mortality) and standard error were calculated for each concentration. A three-parameter sigmoid curve was fitted to survival data from all females in Sigmaplot to assess adult female LC_50_ at 21 days; male LC_50_ could not be calculated as males did not reach 50% mortality at 21 days at any tested mannitol concentration.

Differences in emerging adult sex ratios were tested using a chi square in GraphPad against an expected 50-50 male-female ratio [28], with the same sample size as in the treatment vial. All treatment vial sex ratios were also tested against our control vials using a chi square. Differences in mean time to eclosion (in days) of males and females across concentrations were assessed using a pairwise log-rank Mantel Cox survival analysis test. Linear regressions of male and female eclosion day vs. mannitol concentration, with comparisons of slopes and intercepts, were used to determine if the concentration of mannitol had a similar effect on eclosion day in both sexes.

## Results

### Concentration-Dependent, Sex-Specific Adult Longevity to 21 Days

Flies fed control foods without molasses did not differ in their longevity to 21 days from those fed control foods with molasses (X^2^=0.2, p=0.66), demonstrating that this source of carbohydrates is not necessary for adult *D. melanogaster* survival to 21 days, and that leaving molasses out of the mannitol treatment foods did not affect longevity during this time period.

Longevity over 21 days was dependent on mannitol concentration for both sexes. Female longevity did not significantly differ from the no molasses controls in the 0.25M treatment (X^2^=0.73, p=0.39), but differed significantly in all other mannitol treatments (Figure 1; 0.5M: X^2^=4.30, p=0.04; 0.75M: X^2^=24.84, p<0.001; 1M: X^2^=37.88, p<0.001; 1.5M: X^2^=17.82, p<0.001; 1.5M: X^2^=26.788, p<0.001). Male longevity to 21 days did not differ significantly from the no molasses controls in the 0.25M (no difference observed) or 0.5M (X^2^=3.07, p=0.08) treatment, but differed significantly in the other treatments (Figure1; 0.75M: X^2^=8.81, p=0.003; 1M: X^2^=15.60, p<0.001; 1.5M: X^2^=7.86, p=0.005; and 2M: X^2^=7.69, p=0.006).

**Figure 1.**
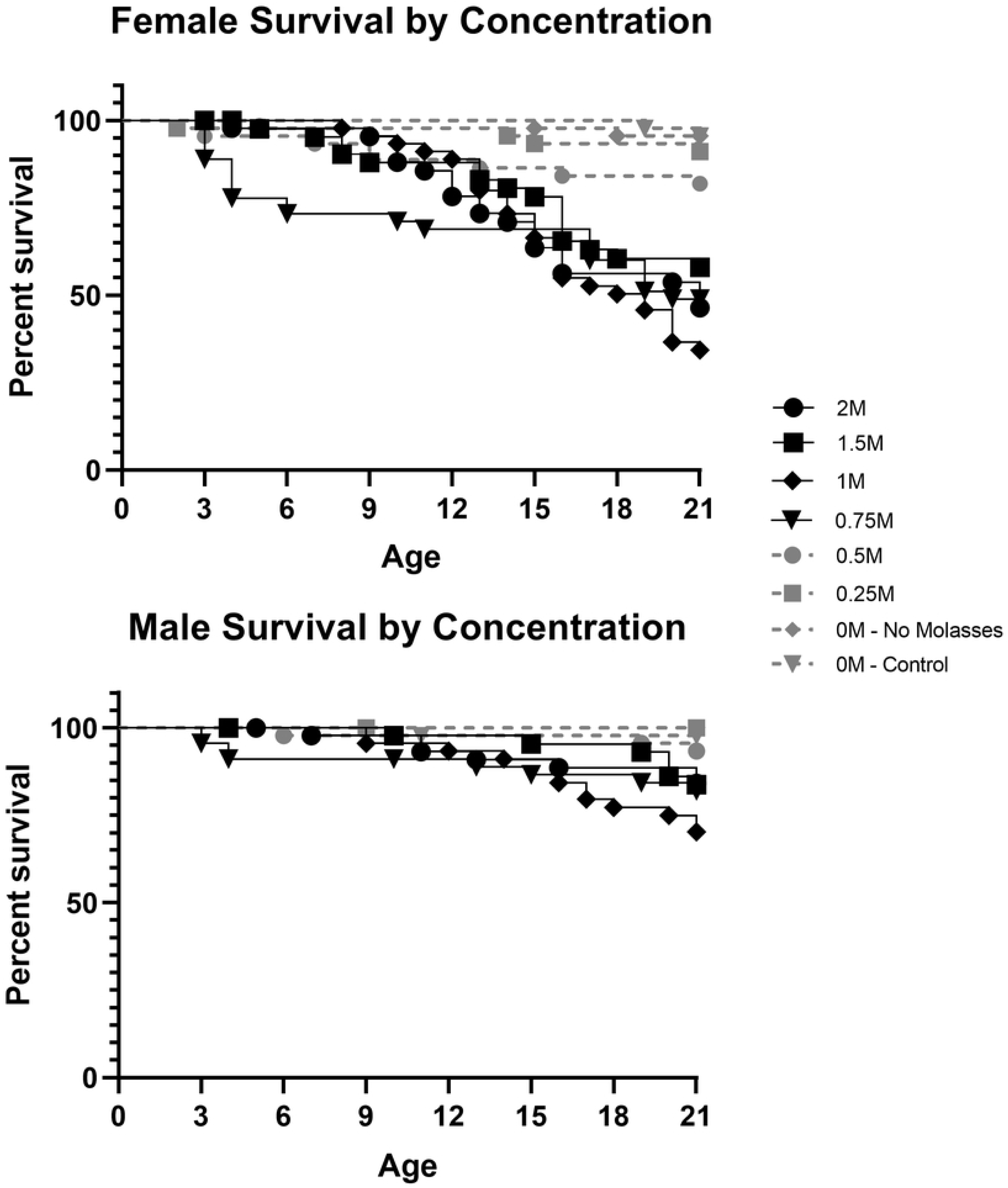
Survival plots showing percent survival versus adult age of female (top) and male (bottom) *D. melanogaster* given control food or foods with increasing concentrations of mannitol (0.25M to 2M). Observations were terminated at 21 days of age (n=45flies/sex/treatment). Highly significant differences (p<0.01) from the no molasses control are in black, non-significant differences are in grey.

The best-fit sigmoidal curve for adult female LC_50_ data at 21 days was:

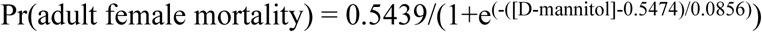

This curve was a significant fit to the data (Figure S1; R^2^=0.898, p=0.013) and using the equation we estimated the adult female LC_50_ at 21 days to be 0.76M mannitol. Males did not have an LC_50_ in this experiment; maximum adult male mortality at 21 days was 30.2%, in the 1M mannitol treatment.

Adult male and female flies did not differ in their longevity to 21 days in either 0M condition (control: X^2^=0.33, p=0.57; no molasses: X^2^=2.02, p=0.16) or the 0.5M mannitol treatment (X^2^=2.79, p=0.09). The sex difference was marginally significant in the 0.25M treatment (X^2^=4.07, p=0.04). Adult females had highly significantly decreased longevity relative to males in the other treatments (Figure 2; 0.75M: X^2^=10.65, p=0.001; 1M: X^2^=12.13, p<0.001; 1.5M: X^2^=7.92, p=0.005; 2M: X^2^=13.54, p<0.001).

**Figure 2.**
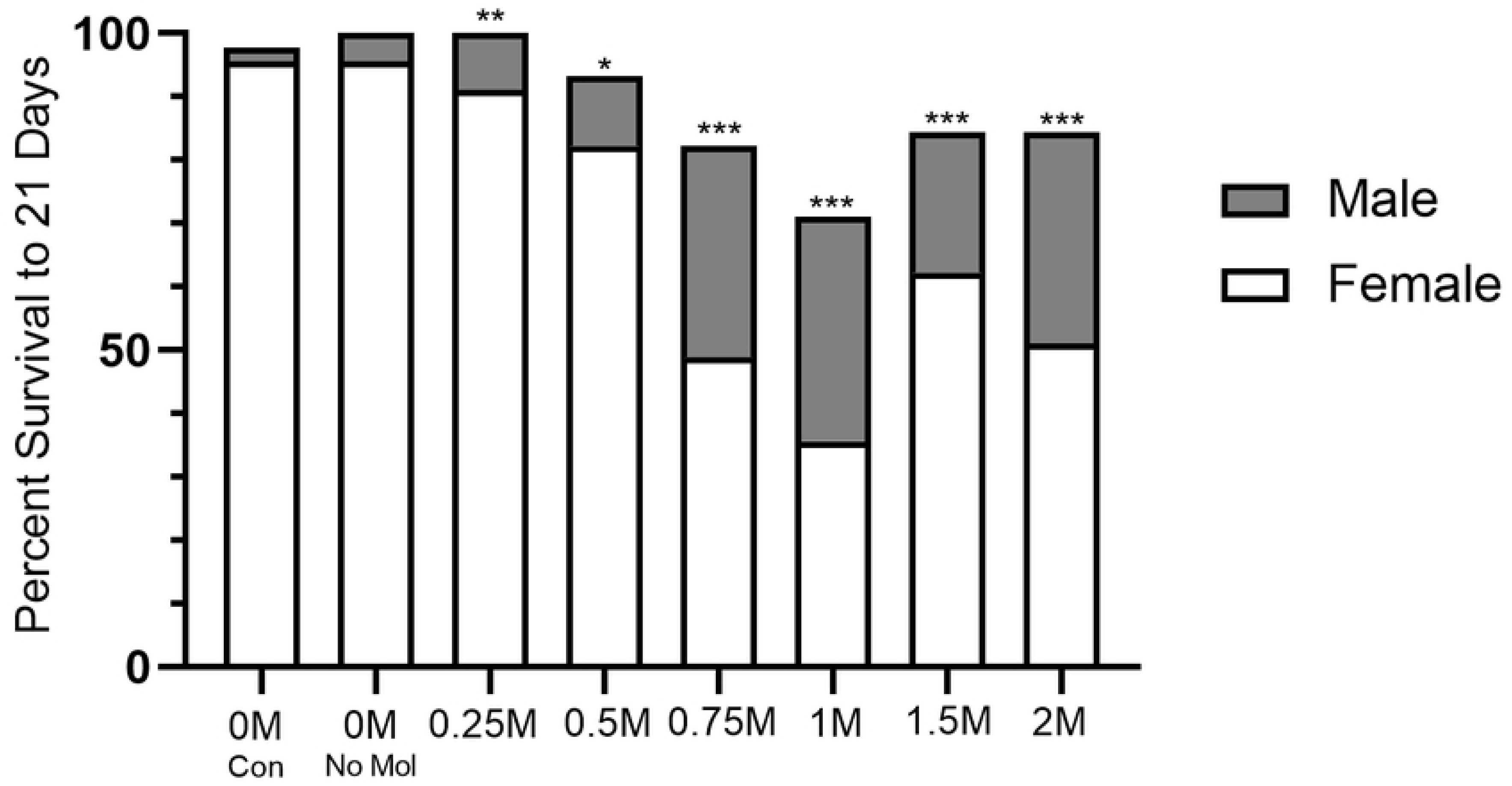
Plot showing the difference in percent survival of male flies versus female flies when given control food or foods with varying concentrations of mannitol (0.25M to 2M). Observations were terminated at 21 days of age (n=45flies/sex/treatment). Highly significant differences in survival distributions (Mantel-Cox; p<0.01) are denoted by three stars, significant differences (p<0.05) are denoted by two stars, nearly significant differences (p<0.1) are denoted by one star, and non-significant differences have no symbols.

When dead females from mannitol treatments were dissected, white matter was found within the crop and often also around the mouth and anus; this was not true of living flies sacrificed from the 0M foods at the same time point (Figure 3). Blue dye was found in nearly 100% of flies by visual inspection on Day 1 of trials, and fecal matter was blue in all treatment vials, proving that the flies consumed the mannitol foods. However, flies in the mannitol treatments were rarely found with blue in their guts on the day of their death, suggesting the dye was excreted.

**Figure 3.**
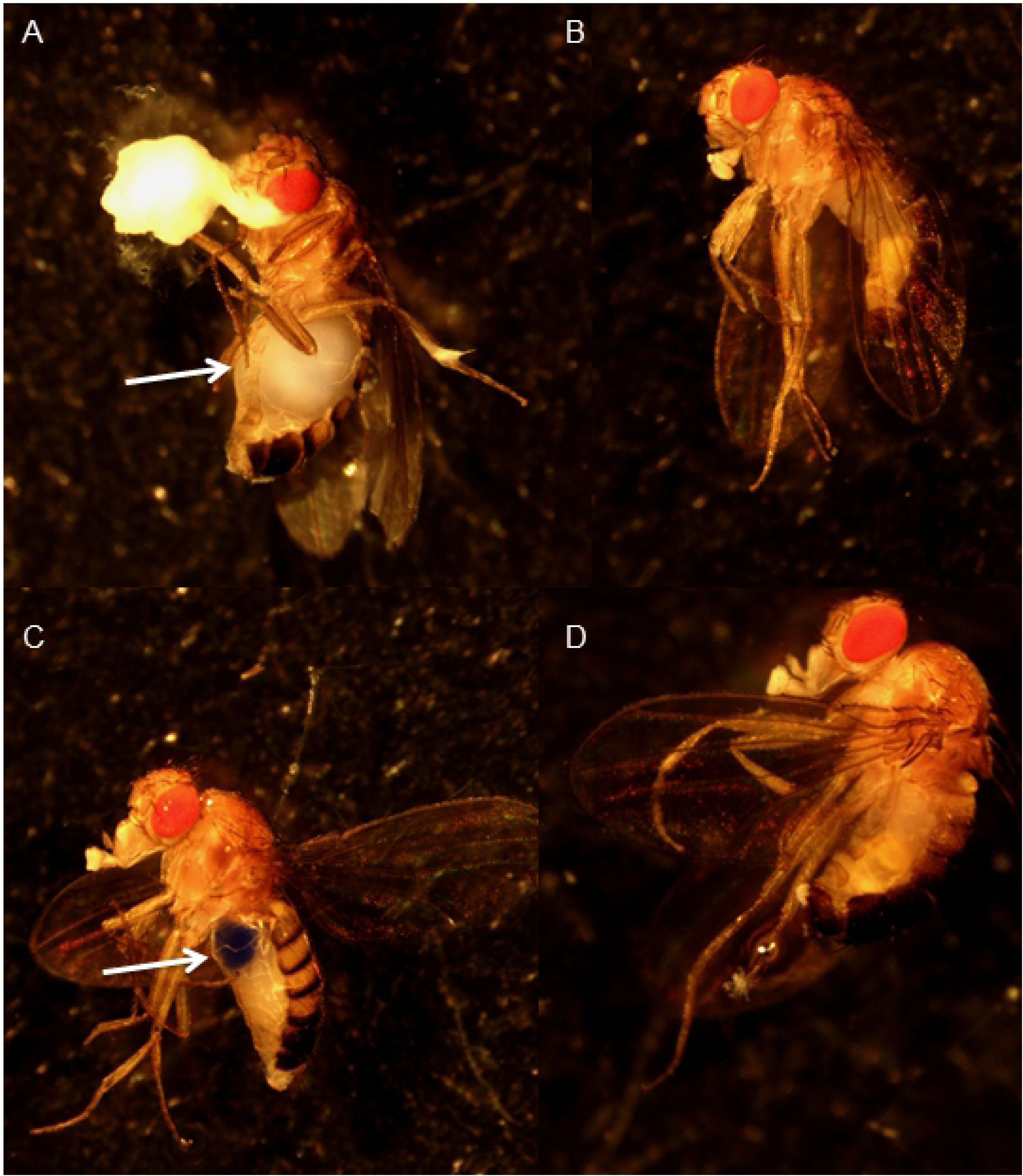
Images of adult CantonS flies taken four days after being set on mannitol foods. A) FeAmale fed 0.75M mannitol and found dead inBtreatment before photographing. Arrow indicating accumulated white mass in the crop. B) Male fly sacrificed for photograph and was alive in 0.75M mannitol treatment. C) Female sacrificed from 0M mannitol food. Arrow indicating blue dye in the crop. D) Male sacrificed for photograph from 0M mannitol treatment.

### Differences in Longevity to 21 Days in Mixed Sex vs. Single Sex Vials

Females flies in mixed-sex vials had significantly reduced longevity to 21 days compared to females kept in single-sex vials in the 1M, 1.5M, and 2M treatments (Figure 4; 1M: X^2^=8.67, p=0.003; 1.5M: X^2^=9.73, p=0.002; 2M: X^2^=4.12, p=0.04) but not in the 0M control, nor in the lower concentration treatments (0M: X^2^=0.28, p=0.6; 0.25M: X^2^=0.46, p=0.5; 0.5M: X^2^=3.33, p=0.07; 0.75M: X^2^=2.83, p=0.09). A similar pattern was observed for males (Figure 4; 1M: X^2^=12.65, p<0.001; 1.5M: X^2^=5.80, p=0.02; 2M: X^2^=5.92, p=0.02; 0M: X^2^=2.00, p=0.16; 0.25M: X^2^=n.d., p=n.d.; 0.5M: X^2^=1.50, p=0.22; 0.75M: X^2^=0.93, p=0.33). In both cases, the majority of the difference in survival between single-sex vials and mixed-sex vials occurred after 12-15 days (Figure S4).

**Figure 4.**
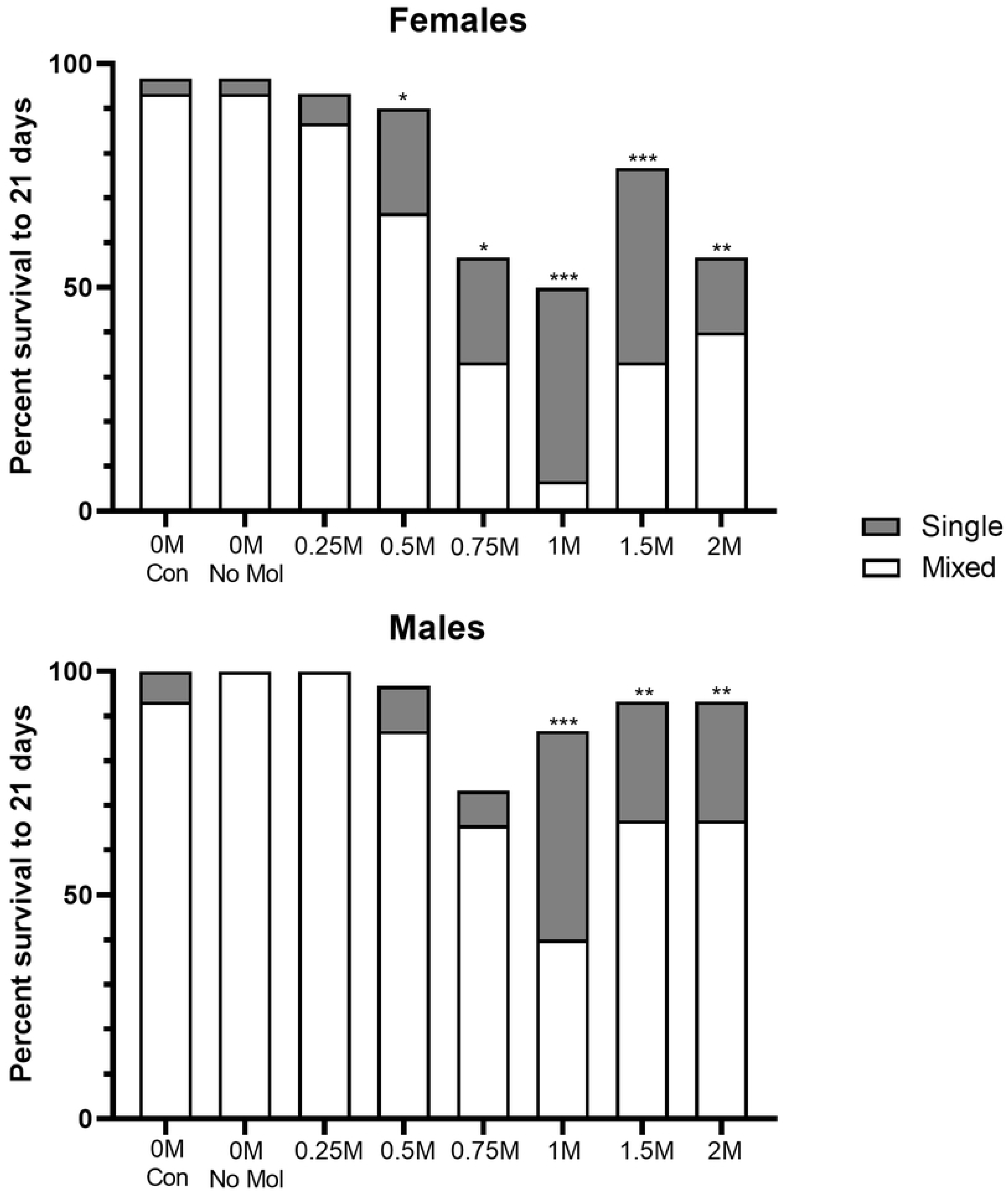
Plot showing the difference in percent survival of flies cultured in single sex vials versus flies of the same sex cultured in mixed sex vials when given foods with the same concentration of mannitol. Observations were terminated at 21 days (n=30 flies/sex for single sex treatments; n=15 flies/sex for mixed-sex treatments). Highly significant differences in survival distributions (Mantel-Cox; p<0.01) are denoted by three stars, significant differences (p<0.05) are denoted by two stars, nearly significant differences (p<0.1) are denoted by one star, and non-significant differences have no symbols.

### Effects of Mannitol Fed to Larvae on Adult Sex Ratios and Eclosion Day

We asked whether larvae fed mannitol would experience the same sex-specific mortality found in adults. Expected adult sex ratios upon eclosion are 1:1 [28]; none of our conditions deviated from the expected ratio (Chi square, 0M: p=0.71; 0.4M: p=0.13; 0.8M: p=0.73), nor did treatment conditions deviate from the sex ratio of our 0M condition (Chi square: p=0.20). Eclosion day was significantly later in both males and females in 0.4M trials compared to controls (Figure S2: Mantel-Cox, males: X^2^=308.51, p<0.001; females: X^2^=491.21, p<0.001) and even more delayed in 0.8M conditions than 0.4M (Mantel-Cox, males: X^2^=401.47, p<0.001; females: X^2^=448.98, p<0.001). Females and males did not differ in how much increasing mannitol concentration delayed their development (Figure S3: slopes: p=0.14; elevations: p<0.0001) but males in all conditions emerged later than females at their concentration (Mantel-Cox, p<0.001).

## Discussion

Mortality induced by mannitol was concentration dependent for both sexes. Female adult flies ingesting mannitol showed significant, concentration-dependent decreases in longevity to 21 days compared to controls from 0.5M-2M mannitol, while males showed significant decreases in longevity compared to controls from 0.75M-2M mannitol. Females had an LC_50_ of 0.76 M at 21 days, while males reached a maximum mortality of 30.2% (in the 1M treatments).

Decreases in adult female longevity to 21 days were greater than male decreases in longevity at concentrations of 0.75M and above. Our results supported our hypothesis that increased food ingestion, and therefore self-dosing with mannitol, due to differing reproductive costs generates sex-specific differences in mortality between males and females. Males and females have differing nutritional requirements generated by the different energetic demands of reproduction [20]. Females feed more frequently, and consume greater volumes of food, than males because oogenesis is both energetically and nutritionally costly [13-14].

Differences in longevity to 21 days between single-sex and mixed-sex vials may also be related to reproductive energetic demands that generate differences in food ingestion and self-dosing between members of the same sex. Single-sex vial females and males were likely to be virgins as they were only housed with the opposite sex for up to 24 hours post-eclosion (period of low receptivity to mating in both sexes) [29-31]. Mating females and males (kept in mixed-sex vials) had significantly decreased longevity compared to virgin females and males (kept in single-sex vials) at concentrations of 1M, 1.5M, and 2M mannitol. Actively mating males and females reproduce more than virgin males and females, and mating and reproduction are energetically costly in both sexes [32-34]. Fertile *D. melanogaster* females feed more than sterile or virgin females and the receipt of sex peptides also induced feeding and reproduction in fertile females [15,35]. *Drosophila bifurca* males raised with infrequent access to females produced significantly less sperm than those in mixed-sex conditions [34]. Behaviors associated with courtship, competition, and mating may also be energetically costly for males, causing increased food ingestion, and thus mannitol self-dosing, for males in mixed-sex vials [18-20,22].

Female longevity decreases with male encounter rates and with exposure to male seminal proteins (sex peptides) [28,36], however females in mixed-sex control vials did not differ in their longevity from females in single-sex control vials. Additionally, male exposure alone would not explain the sharp decrease in male survival also seen in mixed-sex vials at high mannitol concentrations. Social contact has also been posited to decrease longevity in adult flies [21], but neither males nor females showed significant differences in longevity based on culturing condition at lower mannitol concentrations (0.25M to 0.75M). Increased self-dosing of mannitol via increased food ingestion in order to meet the energetic demands of mating and reproduction may cause the mating vs. virgin effect seen for both males and females in this experiment at high concentrations.

Emerging adult sex ratios in vials of larvae fed mannitol did not differ significantly from sex ratios in control vials, or from a 50-50 sex ratio. Both male and female larvae fed mannitol also had similar increases in their mean times to eclosion over increasing mannitol treatments. The delayed eclosion of larvae of both sexes (and lack of a sex-specific mortality in emerging adults) may indicate significant differences in how mannitol impacts developmental stages of the same species. In addition, ∼70% fewer larvae of either sex emerged in the 0.8M as compared to the 0M vials (while 0M and 0.4M saw similar numbers) – however, because we did not control for the number of eggs laid in each vial, we cannot definitively state that mannitol reduced survival in larvae.

Our data showed a significant difference in response to mannitol based on sex in adults only. An alternative hypothesis to reproductive demands is that the presence of mannitol in food signals different behaviors in adult males and females, as sexual dimorphism has been found in *D. melanogaster* neuronal responses to particular carbohydrates [37]. Little research has focused on mannitol perception in *D. melanogaster*, though mannitol receptors have been found in other insect species [8]. Gr5a sugar neurons show a small response to the administration of 100mM mannitol [38]. Some receptors from the Gr64a-f group are also known to modulate neural responses to sugar alcohols and stimulate feeding on fermenting yeast products that may produce mannitol [3,39-41]. These neurons should be further investigated for their ability to modulate neuronal responses to mannitol and to look for dimorphism between the sexes.

Our hypothesis that reproductive energetic demands drive mannitol ingestion and differences in mortality between sexes, and between mating vs. virgin flies, does not explain the mechanism by which mannitol ingestion kills *D. melanogaster* adults. Dead females from 0.75M mannitol treatments had white matter in their crops as early as day four; this accumulation was noted in nearly all dead flies of both sexes across concentrations when dissected. This suggests that some food components, including the blue dye, were excreted by the flies while others remained and accumulated in the gut. If mannitol cannot be metabolized efficiently it may accumulate in the crop. *T. castaneum* beetles likely utilizes NADP^+^-dependent D-arabitol dehydrogenase for mannitol catalysis [9] but this enzyme is not found in *D. melanogaster* [42]. Sorbitol dehydrogenase (SDH) is another possible pathway for mannitol breakdown [43], as mannitol is an isomer of sorbitol [44] and SDH is found in *Drosophila* (Gene ID: 40836, 41313) [45]. Overloading of this pathway has been shown to lead to sorbitol accumulation in the gut [46] and the rate of mannitol catalysis by SDH is very low [43,47]. Taken together, *D. melanogaster* and its common microbes may be able to metabolize small amounts of mannitol, but likely not the amounts consumed in our experiment [48-51]. Mannitol may also slow down digestion [6], elevate carbohydrate ingestion to a detrimental degree [52], or have a dieuretic effect due to its slow absorption in the gut [53-54]. In flies, it is possible mannitol is transported into the hemolymph by aquaporins from the midgut [55], where it may lethally increase osmotic pressure like other sugar alcohols [56-57].

Future work should attempt to pinpoint the mechanism of mannitol’s lethality in *D. melanogaster*, particularly in comparison to other insect taxa where it is nutritive. Possible differences in feeding behavior between males and females could be caused by differences in mannitol perception that should be explored. Additionally, mannitol seems to directly impact the timing of key developmental stages pre-eclosion. The effects of mannitol on larval and juvenile stages have not been studied in any insect species to date, but we have demonstrated differences in mannitol’s effects across developmental stages.

## Conclusions

Mannitol caused concentration-dependent decreases in longevity to 21 days in both males and females at concentrations above 0.75M (males) or 0.5M (females). Female longevity was more significantly decreased than male longevity at concentrations above 0.75M. Actively mating males and females had decreased longevity compared to virgin males and females at concentrations of 1M, 1.5M, and 2M. Mannitol fed to larvae did not alter emerging adult sex ratios, suggesting that sex-specific mortality due to mannitol occurs only in adults. Overall, our results support our hypothesis that sex differences in energy needs, related to the nutritional and behavioral demands of mating and reproduction, contribute to decreased longevity in females compared to males. We further conclude that both males and females have mannitol-induced decreases in longevity when mated, as compared to virgins, due to the increased costs of reproduction for both sexes.

## Acknowledgements

We thank Angelina Gomez, Natalie Carroll, Virginia Caponera, and Devneet Kainth for assistance caring for and counting adults. Kaitlin Baudier provided early assistance with basic methodological and statistical procedures. Jennifer Viveiros provided support and training in fly husbandry, sexing, and making foods. Funds from Drexel University Office of Research supported this work (to DRM). The funders had no role in study design, data collection and analysis, decision to publish, or preparation of the manuscript. Some of this material is based upon work supported by (while serving at) the National Science Foundation (for DRM). Any opinion, findings, and conclusions or recommendations expressed in this material are those of the author(s) and do not necessarily reflect the views of the National Science Foundation.

## Supplementary Figures

**Figure S1.** Percent mortality of adult female flies plotted against concentration of mannitol provided in foods. The three-parameter best-fit sigmoidal function is shown and was used to calculate the LC_50_ for flies at 21 days (0.76M mannitol). Error bars represent one standard deviation.

**Figure S2.** Average eclosion day of male and female flies across increasing concentrations of mannitol (0M to 0.8M). Letters indicate highly significant differences (Mantel-Cox, p<0.001) between flies of the same sex. Error bars represent one standard deviation (n=1,249 females; 1,181 males).

**Figure S3.** Linear regressions for male (grey circle) and female (black square) larvae showing the effect of increasing mannitol concentration on eclosion day. Both females and males eclose later when fed increasing concentrations of mannitol. Females: y=4.034x+10.80 (F=1620, R^2^=0.5650, p<0.0001), males: y=4.271x+11.06 (F=1168, R^2^=0.4977, p<0.0001). The slopes of the lines are not significantly different (F=2.206, p=0.1376) but the intercepts are (F=68.38, p<0.0001). Error bars represent one standard deviation (n=1,249 females, 1,181 males).

**Figure S4.** Survival plot showing the significant difference in the percent survival of flies in single sex vials over mixed sex vials when given foods with the same concentration of D-mannitol. Observations were terminated at 21 days of age (n=30 flies/sex for single sex treatments; n=15 flies/sex for mixed-sex treatments). Males and females housed together in 1-2M mannitol treatments had much lower survival to 21 days than males or females housed in single-sex vials.

